# Sweeping genomic remodeling through repeated selection of alternatively adapted haplotypes occurs in the first decades after marine stickleback colonize new freshwater ponds

**DOI:** 10.1101/191627

**Authors:** Susan Bassham, Julian Catchen, Emily Lescak, Frank A. von Hippel, William A. Cresko

**Affiliations:** Institute of Ecology and Evolution, University of Oregon, Eugene, OR, 97403, USA; Department of Animal Biology, University of Illinois at Urbana-Champaign, Urbana, IL, 61801, USA; Department of Biological Sciences, University of Alaska Anchorage, Anchorage AK 99508, USA; College of Fisheries and Ocean Science, University of Alaska Fairbanks, Fairbanks AK 99775, USA; Present address: Department of Biological Sciences & Center for Bioengineering Innovation, Northern Arizona University, Flagstaff, AZ, 86011, USA

**Keywords:** contemporary evolution, ecological divergence, population genomics, Gasterosteus aculeatus, threespine stickleback

## Abstract

Heterogeneous genetic divergence can accumulate across the genome when populations adapt to different habitats while still exchanging alleles. How long does diversification take and how much of the genome is affected? When divergence occurs in parallel from standing genetic variation, how often are the same haplotypes used? We explore these questions using RAD-seq genotyping data, and show that broad-scale genomic re-patterning, fueled by standing variation, can emerge in just dozens of generations in replicate natural populations of threespine stickleback fish (*Gasterosteus aculeatus*). After the catastrophic 1964 Alaskan earthquake, marine stickleback colonized newly created ponds on seismically uplifted islands. We find that freshwater fish in these young ponds differ from their marine ancestors across the same genomic segments previously shown to have diverged in much older lake populations. Outside of these core divergent regions the genome shows no population structure across the ocean-freshwater divide, consistent with strong local selection acting in alternative environments on stickleback populations still connected by significant gene flow.Reinforcing this inference, a majority of divergent haplotypes that are at high frequency in ponds are shared across independent freshwater populations and are detectable, at low frequency, in the sea even across great geographic distances. Building upon previous work in this model species for population genomics, our data suggest that a long history of divergent selection and gene flow across stickleback in oceanic and freshwater habitats has created balanced polymorphism in large genomic blocks of alternatively adapted DNA sequences, ultimately stoking - and potentially channeling - rapid, parallel evolution.

## Introduction

When populations of a species adapt to different environments or habitats, parts of the genome may chart divergent evolutionary courses, leading to heterogeneous patterns of genomic differentiation (1-3). Recent research demonstrates that complete genetic isolation is not a prerequisite for such divergence (4-6). While these genomic patterns can occur neutrally via genetic drift in isolated populations, pronounced heterogeneous genomic divergence might often be shaped by the action of local adaptation to different habitats combined with gene flow among the populations (7, 8). Under strong natural selection, alleles and genotypes that are alternatively adapted to different habitats can be maintained at divergent frequencies even when the populations are still exchanging alleles. Local adaptation and gene flow can result in differentiation at selected loci and homogenization of variation in neutral regions of the genome (8, 9). When such divergence with gene flow plays out over geographic space, the metapopulation structure of a species can promote the maintenance and reuse of standing genetic variation for adaptation and even speciation (10-12).

Cases of rapid adaptation (13-16) - so-called contemporary evolution (17-21) - often invoke the use of standing variation because the extended time needed to accumulate new beneficial mutations is presumed to limit the rate of adaptation (22-25). Although standing genetic variation permits accelerated change, the patterns of heterogeneous genomic differentiation that materialize during contemporary evolution still remain largely unexplored in natural systems (26) particularly in the context of local adaptation with gene flow (27). It is still not known what proportion of the genome is affected during contemporary evolution by direct selection on standing adaptive variants, and how much of the genome is influenced by linked selection such as genetic hitchhiking and genetic draft (28-31). When a species exists across diverse habitat types with distinct selective pressures, to what extent do independent populations adapting to similar habitats initially make use of the same haplotypes, and how and at what frequencies are these adaptive haplotypes distributed and maintained species-wide? A first step in tackling these questions is to survey heterogeneous genomic re-patterning using haplotype-level data in organisms while they are first evolving in response to different natural habitats. Across the northern hemisphere marine threespine stickleback fish (*Gasterosteus aculeatus*) have invaded and adapted to countless freshwater habitats over multiple time scales, making them a fruitful model for vertebrate evolution (32-34) and for population genomics of rapid adaptation (35). Marine and freshwater stickleback differ considerably in many traits (36-40), and remarkably parallel phenotypic and genomic evolution has followed independent invasions by marine fish into freshwater habitats, including those on landscapes de-glaciated only thousands of years ago[41-49]. At least some of this phenomenon might be due to the reuse of standing genetic variation (41-43), enabling the rapid and parallel adaptive evolution documented in this species (44-51). Schluter and Conte (26) framed one possible scenario to explain these dynamics: a freshwater-adapted stickleback genotype, scattered into the marine population by gene flow and recombination, might be brought together and reintegrated during subsequent colonizations of freshwater habitats. What is still needed in stickleback is investigation of the time scale on which reassembly can transpire, which depends strongly on the frequencies and genomic granularity of freshwater adapted haplotypes carried by marine populations. Both of these qualities have been largely unquantified.

Here we explore the nature of this ‘freshwater genome scattering and reassembly’ during the earliest stages of freshwater adaptation by marine stickleback, using naturally formed stickleback populations for which convincing genetic and geologic evidence indicates they colonized new freshwater habitats within the last 50 years (35). Submarine terraces that encircled islands in the Gulf of Alaska and Prince William Sound were suddenly thrust above sea level in 1964 by the Great Alaska Earthquake (52-54), and the resulting changes in sediment deposition and erosion over the following years created new freshwater ponds (35). Threespine stickleback fish that now inhabit many of these ponds have become as morphologically distinct in degree and form from their immediate marine ancestors as have much older mainland Alaska freshwater stickleback populations (35, 54, 55), similar to phenomena documented in older freshwater populations throughout the northern hemisphere (33, 36, 56-58).Previously, using a relatively small number of randomly chosen genetic markers in these new populations, we showed that marine fish likely invaded and adapted to fresh water several independent times on different islands and even among ponds on the same island (35).

The present study wields a much denser dataset of loci in over 1200 fish from independently derived stickleback populations on three seismically uplifted marine islands (Danger, Middleton and Montague Island), as well as from mainland Alaska and Oregon. We use this dataset to dissect genome-wide patterns of haplotype frequency divergence across populations that range in age from just decades to thousands of years. By exploring these replicate natural experiments in adaptation to alternative environments, we find that a much larger proportion of the genome is affected - through combined effects of direct and linked selection - than had been previously reported in stickleback. Loci with alternative haplotypes that dominate in either marine or freshwater populations are clustered in the genome, particularly across broad regions of divergence where marine fish are relatively depleted in haplotype diversity. We show how marine stickleback, even at great geographic distances, act as carriers of a common set of freshwater genotypes, leading to repeated selection of the same haplotypes in independently colonized lakes and ponds.

## Results

### Parallel patterns of genomic divergence involving a quarter of the genome assemble in decades on a background of no population structure

Despite their young age and their isolation by ocean expanses, we find that stickleback populations newly adapted to fresh water on seismically uplifted Middleton and Montague Islands (south of mainland Alaska) reiterate genomic patterns first described in fish from much older, post-glacial lakes on the Alaskan mainland (46) (Fig 1). We calculated genome-wide population genetic differentiation statistics (F_ST_, F_ST_´, and ϕST; Table S1) using RAD-seq genotyping in stickleback sampled from multiple populations, and we performed genome scans using continuous divergence values (Fig 1; Fig 2).

**Fig 1.**
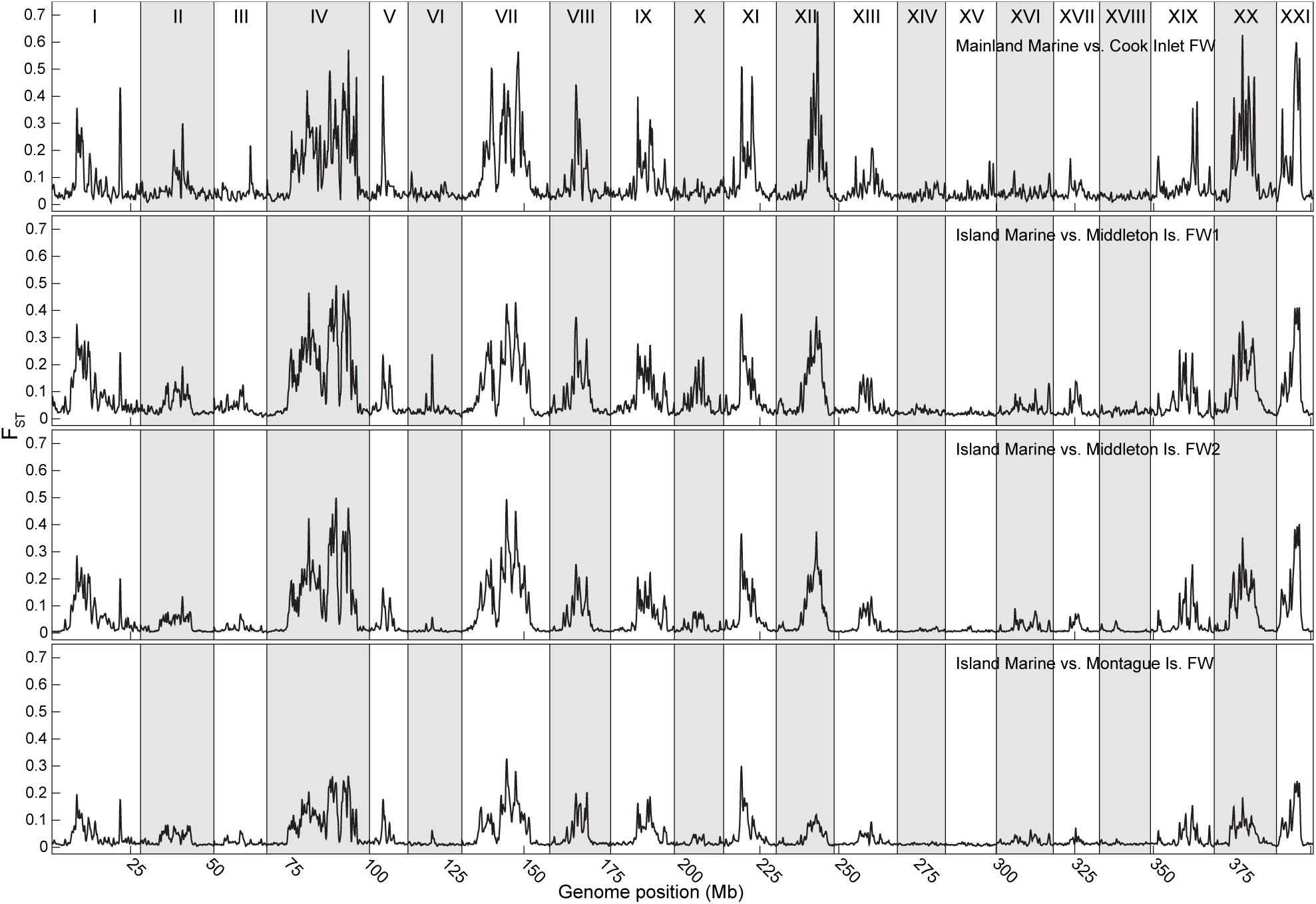
Nascent and old freshwater stickleback populations share parallel patterns of genomic divergence from marine fish. Smoothed, SNP-based FST is plotted along the genome in comparisons of mainland coastal marine versus mainland post-glacial Cook Inlet freshwater, and island marine fish (from collection sites Da02, Mi17, Mi23) versus freshwater populations that formed after the 1964 earthquake on geographically separated islands: Middleton Island (FW1: Mi07, Mi17; FW2: Mi06, Mi11, Mi16, Mi28) and Montague Island (Mo30, Mo33). These populations were found to form cohesive groups via genetic structure analyses in Lescak et al. [35]

**Fig 2.**
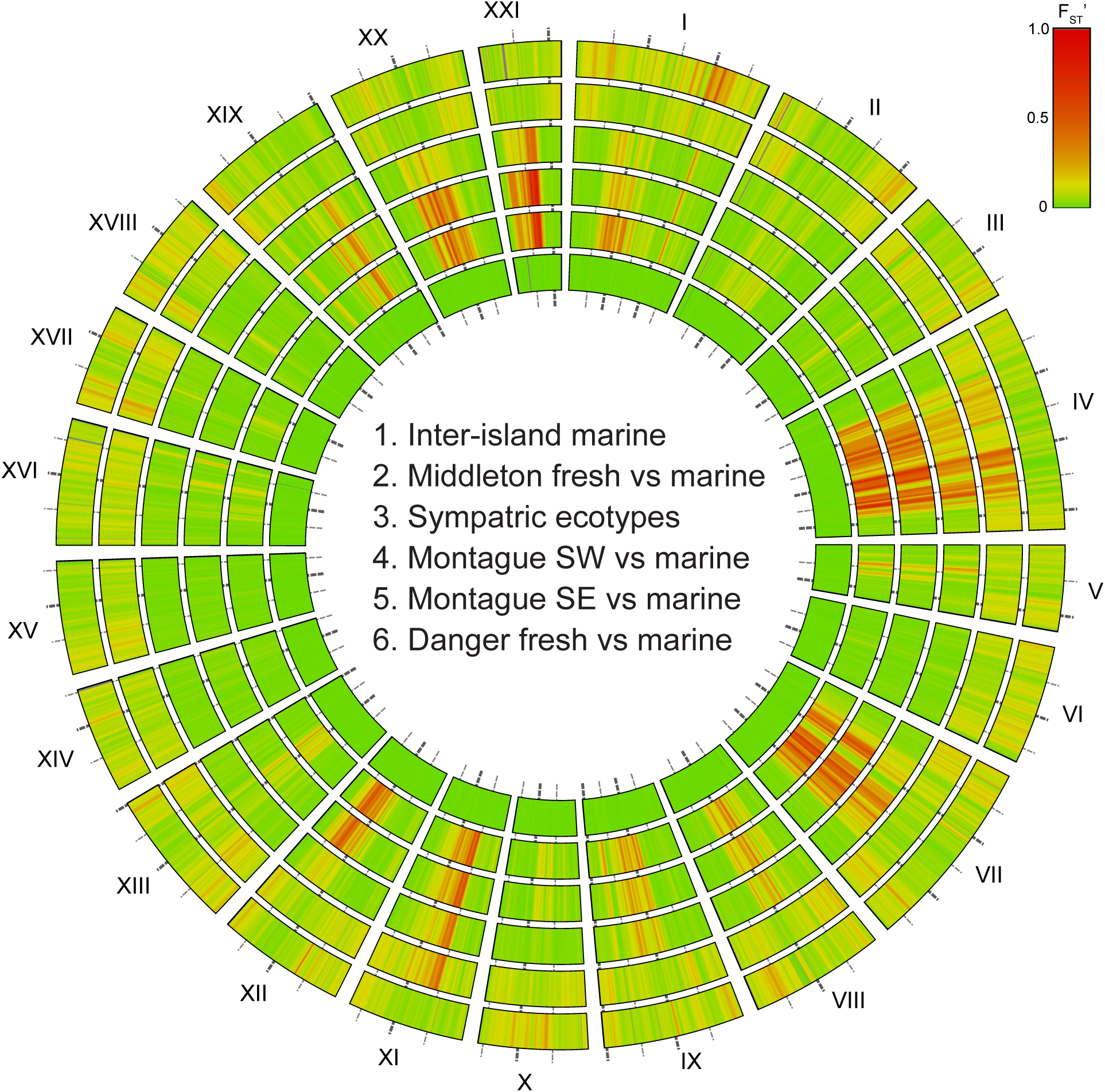
Patterns of genomic divergence are similar among young, independently colonized freshwater locales. Smoothed, haplotype-based FST´ is plotted along the genome. Concentric rings are numbered from the center out, and labeled with 5 Mb intervals. Marine populations (Mi17 vs. Da02) from different islands about 100 km apart show little differentiation (Ring 1). While Middleton Island (MiFW2, Ring 2), sympatric freshwater vs. marine ecotypes both from Mi08 (Ring 3), and SW Montague Island (Ring 4) all show parallel patterns of marine-freshwater divergence, the likely younger populations on Montague (SE, Ring 5) and Danger Island (Da04, Ring 6) differ strongly from marine on only a subset of the typically divergent linkage groups.

A history of multiple founding events from the sea is reflected in the clear population structure that we documented previously (35) among groups of freshwater ponds on three different seismically uplifted islands (Fig S1). Using these pre-defined population genetic groupings for our analyses, we found that the genomic patterns of divergence from marine genomes in one Montague Island group (MoSW) and two groups on Middleton (MiFW1, MiFW2), despite their independent colonization, are broadly congruent, resembling patterns typical of much older populations (Fig 1; Fig 2). A striking pattern is that, outside of foci of high divergence, baseline genetic divergence approaches zero between marine fish and freshwater fish in the new ponds. In southeast Montague (MoSE) and Danger Is. populations, though, we detected only a subset of the F_ST_´ signatures of selection common in freshwater-adapted fish (Fig 2).

In order to quantify what proportion of the genome has diverged over such a short time frame, and to what degree these regions of divergence are shared across independently evolved populations (Fig 1; Fig 2), we developed a Hidden Markov Model (HMM). The emission and transition parameters of the HMM were trained on patterns of F_ST_´ values for RAD loci across the genome in freshwater-marine comparisons. The trained model delineated two states: diverged or not diverged (Fig 3; Fig S2). We found that a significant proportion of the genome falls within well-defined regions that diverge between marine and fresh water in all comparisons: 23% for Middleton FW1, 29% for Middleton FW2, 24% for SW Montague, and 18% for a mainland group (Table 1; Table S2). As expected from the conspicuously parallel genomic divergence across comparisons (Fig 1; Fig 2), the genomic blocks are largely nested when compared among all four freshwater groups. The divergent regions of the two Middleton freshwater groups overlap by over 73%, Middleton and SW Montague overlap by over 70%, and these island populations overlap by between 51 and 59% with the much older mainland populations (Table 1; Table S2). Clearly, the majority of the patterns of genomic divergence are shared across populations, young and old, and this shared, persistent divergence amounts to nearly 15% of the genome.

**Table 1.**
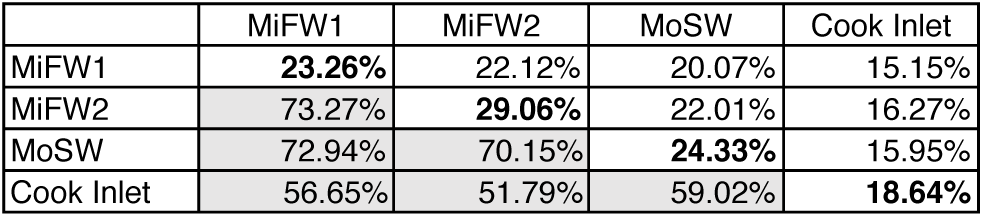
Regions of the stickleback genome that diverge because of adaptation highly overlap between freshwater populations, both young and old. The total amount of the genome allocated to the FW vs. OC “diverged” state by a hidden Markov Model is listed, along the diagonal in bold, for each population. Above the diagonal are the proportions of the genome populations have in common. Below the diagonal is the degree of overlap of the divergent stretches of the genome for each population comparison. Despite the much older time since founding, the Cook Inlet freshwater group still shares more than 50% of divergent regions with the much younger uplift populations.

**Fig 3.**
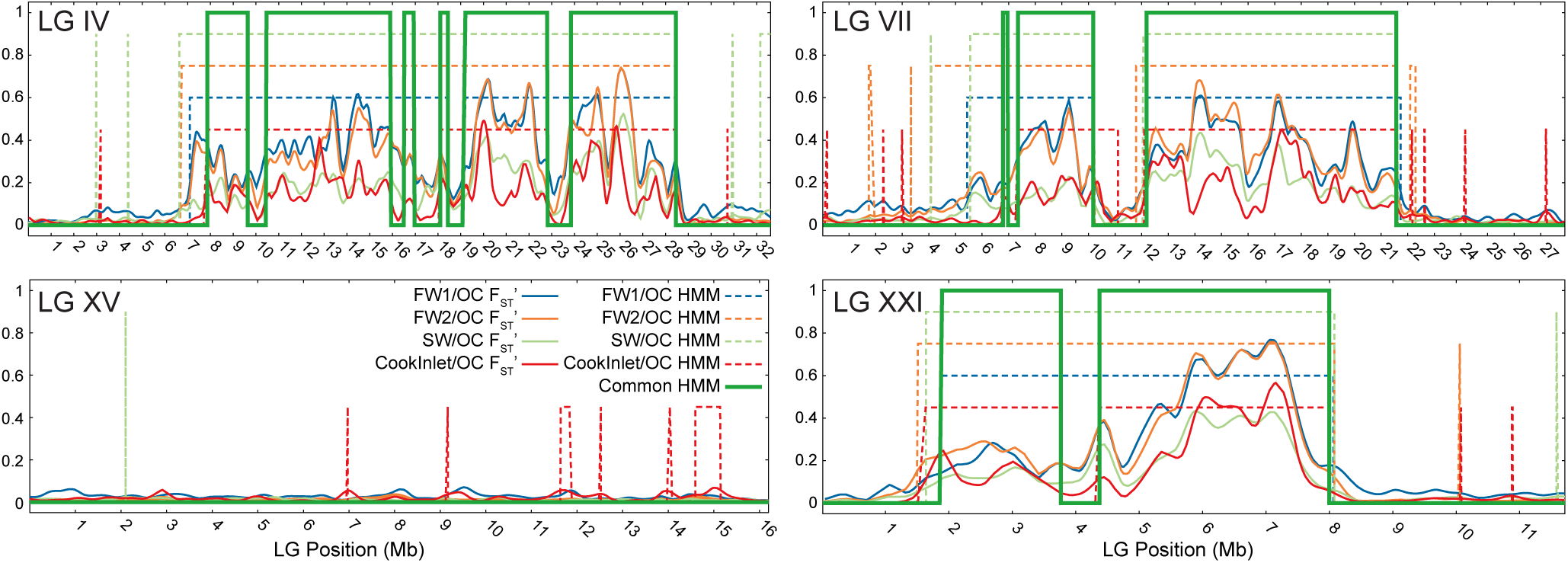
Genomic blocks of freshwater-marine divergence largely overlap among freshwater populations. A) A Hidden Markov Model was trained on FST´ patterns to discriminate between diverged and not diverged regions. Shown here, for example, is the outcome of the model on three linkage groups that have large regions of divergence and one linkage group that has little divergence. From this, the proportion of the genome diverged in each freshwater group (dashed lines) and the divergence overlap (bold green line) among these populations were calculated. B) The total proportion of the genome diverged from marine in the compared freshwater populations ranges nearly to 30%, the majority of which (totaling ≈15% of the genome) overlaps across freshwater populations.

### Marine and freshwater populations differ in patterns of haplotype diversity and composition

F_ST_ statistics quantify relative divergence affected by differences in genetic frequencies among, as well as diversities within, populations (59-61). Because it generates DNA sequence data, RAD-seq allowed us toexamine both the relative and absolute haplotype diversities within differentiated genomic regions in each population, as well as document the distribution and abundance of particular haplotypes across populations. We found that divergent genome segments correspond to regions where marine and freshwater populations differ in haplotype diversity. Marine genomes have lower absolute haplotype diversity across these regions, relative to non-divergent parts of the genome (Fig 4; Fig S3).

**Fig 4.**
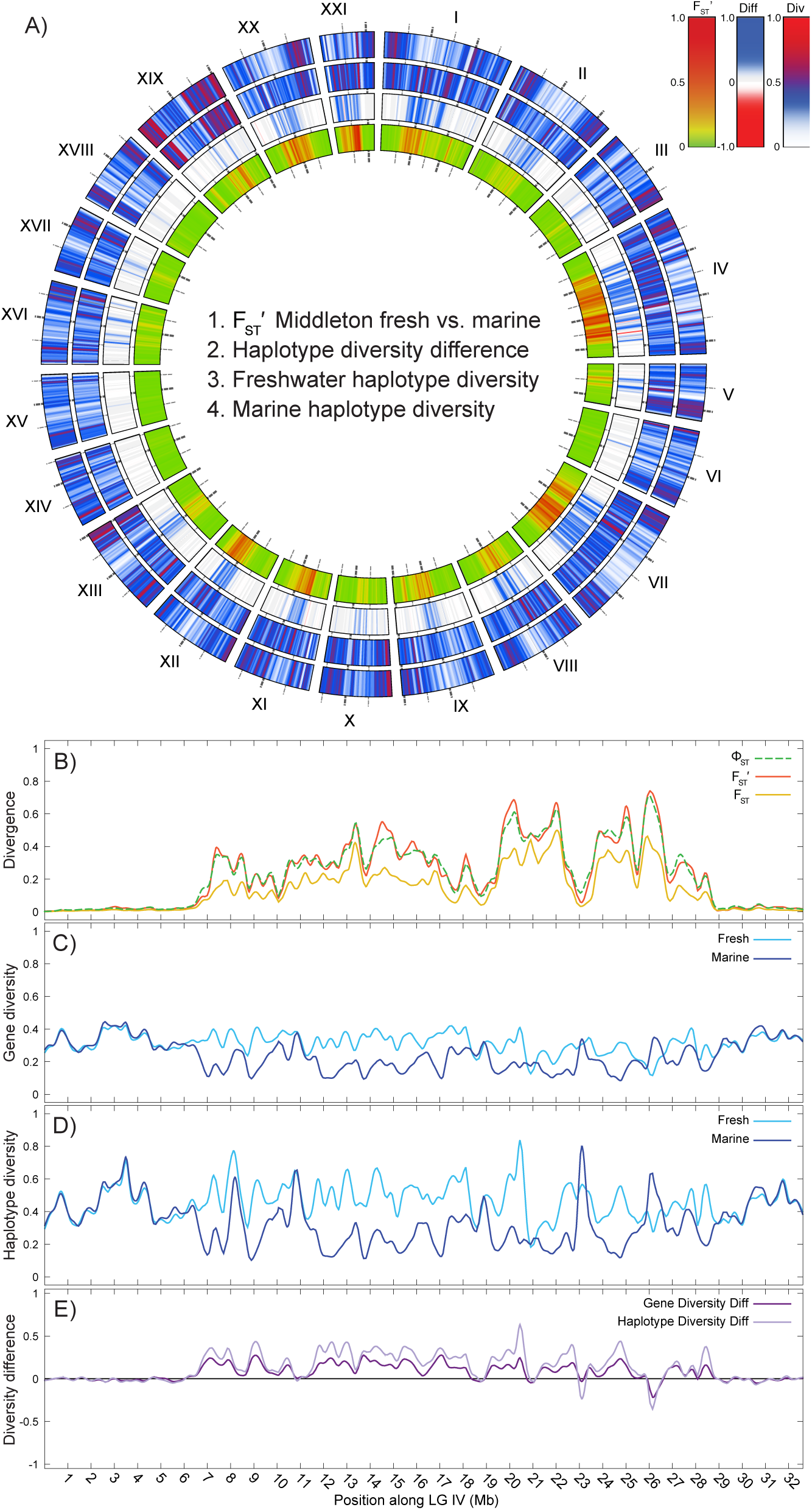
Divergent genomic regions differ also in haplotype diversity between freshwater and marine fish. A) Concentric rings are numbered from the center out. Blocks of elevated FST´ in a comparison of Middleton FW2 vs. marine (Ring 1) highly correspond with relative increases in haplotype diversity in fresh water (Ring 2). On Ring 2, blue represents higher diversity in fresh water than in marine, and red is higher diversity in marine than fresh water. Smoothed haplotype diversity is plotted separately for FW2 (Ring 3) and marine (Ring 4). While a fraction of haplotype diversity outliers does exceed the value range shown, > 99.9% of the smoothed values were 1.0 or less. See Methods for how diversities and difference are calculated. B-D) Middleton FW2 and marine are compared across LG IV, a linkage group to which many morphological traits map that differ between marine and freshwater ecotypes [38]: B) divergence statistics, C) gene diversity, D) haplotype diversity, and E) gene and haplotype diversity difference. Where values in panel D are greater than zero, fresh water has greater diversity than marine.

At individual RAD loci with high estimates of F_ST_´ both the marine and freshwater populations each often have a single but different majority haplotype (Table 2; Table S3), where we define majority haplotypes as the highest frequency haplotypes that - singly or in combination - account for at least 60% of genotypes at a given locus (Fig 5). Independent freshwater populations most often have the same majority haplotype across all shared loci at which there is a single, alternative majority freshwater haplotype (which we call “Case 1 loci“; see Fig 5 for definitions of four qualitatively different cases). For instance, 98.8% (1036 of 1052) of the Case 1 loci shared by the two freshwater population groups on Middleton Island have the same majority freshwater haplotype, and of those shared by both MiFW1 and Montague SW, 99.2% (396 of 399) of them have the same majority haplotype.In MiFW1, MiFW2 and MoSW, loci with majority haplotypes alternative to those in OC (referring to samples combined from marine populations Mi17, Mi23 and Da02) are clustered in the divergent regions delineated by the FST´-based HMM (Fig 5). Outside of these divergent regions, loci with freshwater vs. OC alternative haplotypes occur, but the density of them differs by population group, with MoSW having the fewest, MiFW1 having more than MiFW2, and MoSE have the most (Fig 5; Fig S4).

**Table 2.** Freshwater versus marine divergent loci categorized by haplotype characteristics. Loci with FST´≥ 0.2 are tallied for each population group. Of these, the number that fall into each Case (as defined in Fig 5) is given. Also listed, where applicable, is the proportion (“% Rare“) of these loci carrying a freshwater majority haplotype that occurs only rarely (i.e., less than 10% allele frequency) in the sea. For Case 1 is given the proportion (“% Strict Alt“) of loci that satisfy these criteria: they carry a single majority haplotype in freshwater that is rare in the sea and they carry a different single majority haplotype in the sea. The island populations with a pattern of divergence that closely parallels older mainland populations have the largest proportion these of loci with radically alternative FW vs. OC haplotype frequencies.at which there is a single, alternative majority freshwater haplotype (which we call “Case 1 loci“; see Fig 5 for definitions of four qualitatively different cases). For instance, 98.8% (1036 of 1052) of the Case 1 loci shared by the two freshwater population groups on Middleton Island have the same majority freshwater haplotype, and of those shared by both MiFW1 and Montague SW, 99.2% (396 of 399) of them have the same majority haplotype. In MiFW1, MiFW2 and MoSW, loci with majority haplotypes alternative to those in OC (referring to samples combined from marine populations Mi17, Mi23 and Da02) are clustered in the divergent regions delineated by the F_ST_´-based HMM (Fig 5). Outside of these divergent regions, loci with freshwater vs. OC alternative haplotypes occur, but the density of them differs by population group, with MoSW having the fewest, MiFW1 having more than MiFW2, and MoSE have the most (Fig 5; Fig S4).

| Population | Total loci | Case 1 | % Rare | % Strict Alt | Case 2 | % Rare | Case 3 | % Rare | Case 4 |
| --- | --- | --- | --- | --- | --- | --- | --- | --- | --- |
| MiFW1 | 4847 | 2034 | 74.09 | 66.67 | 1800 | 82.83 | 363 | 93.94 | 610 |
| MiFW2 | 3021 | 1294 | 84 | 75.73 | 1230 | 91.14 | 286 | 98.25 | 192 |
| MoSW | 2114 | 494 | 75.51 | 70.24 | 1408 | 90.63 | 87 | 93.1 | 108 |
| MoSE | 4370 | 1121 | 38.72 | 29.17 | 1703 | 73.69 | 261 | 89.27 | 1245 |
| Da04 | 3800 | 1063 | 29.44 | 19.47 | 1249 | 67.17 | 188 | 85.11 | 1265 |

**Fig 5.**
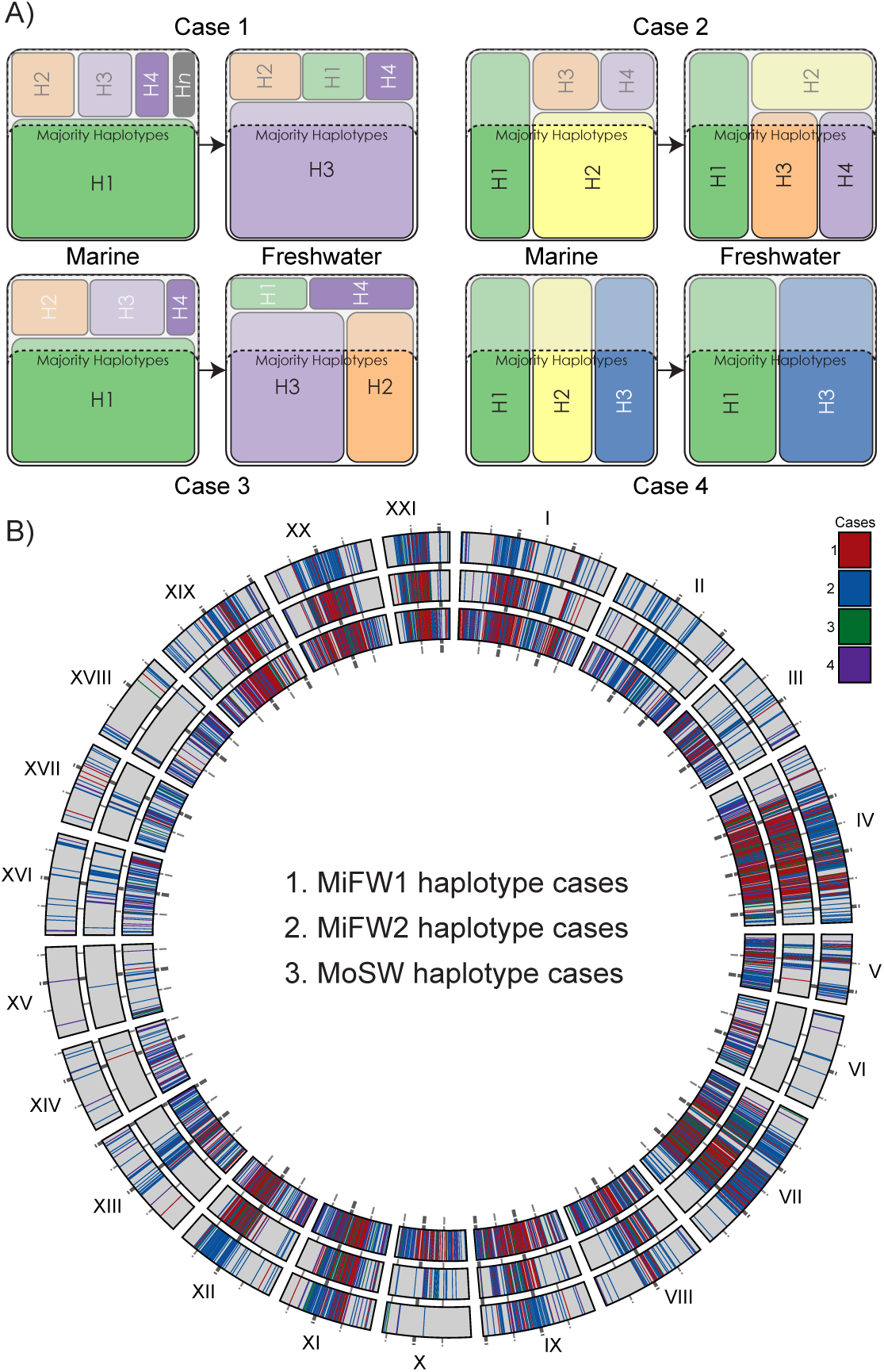
Regions of parallel divergence contain loci with alternative FW versus OC haplo-types. A) To characterize the nature of loci with high FST´ in OC-FW comparisons, we assign these loci to several categories: at Case 1 loci OC and FW populations have single but alternative majority haplotypes (i.e., accounting for at least 60% of population haplo-types); at Case 2 loci, the FW population has one or more haplotypes that are majority (accounting singly or in sum for at least 60% of haplotypes) but are not majority in the sea, and they have at least one majority haplotype that is also a majority haplotype in the sea; at Case 3 loci, the FW population has more than one majority haplotype that is not majority in the sea and no majority haplotypes in common with the sea; and at Case 4 loci, the FW population has majority haplotypes that are a subset of the majority haplotypes in the sea. B) Loci of FST´ ≥ 0.2 are assigned to cases and plotted across the genome for two FW groups from Middleton Is. FW and one Montague Is. While these cases are not strictly discreet, we do think they broadly capture qualitatively different patterns of population genomic processes. Sampling effects could move some loci with borderline allele frequencies (i.e., those with frequencies near our parameter boundaries) between Cases 1-3, but are less likely to move such loci to Case 4 (for example, see Table S3).

Another important qualitative difference between MoSW and Middleton can be seen in the loci falling within the divergent regions; MoSW retains more of the common marine haplotypes at these loci and therefore has fewer loci with only freshwater haplotypes in the majority (i.e., more Case 2 loci, plotted in blue in Fig 5). Taken together, these two distinctions might be expected if MoSW has greater introgression between freshwater and marine fish, as was inferred by Lescak *et al.* (35), who described morphological intermediates in this population. MoSE (which lacks most of the peaks of high F_ST_´) has higher F_ST_´ relative to OC than do the other pond populations across genomic regions that do not usually diverge in fresh water (Fig 2).This pattern is likely a consequence of the low overall haplotype diversity in MoSE, including in regions that house relatively higher haplotype diversity in the sea, perhaps due to small founding population size (Table 3; Fig S3). An interesting correlate of this pattern in MoSE is that F_ST_´ relative to OCdrops across regions normally of high F_ST_´ in other freshwater-marine comparisons because these genomic segments have low diversity in *both* MoSE andOC, and the diffuse number ofhigher F_ST_´ loci found in MoSE predominantly result from subsampling of the majority OC haplotypes (such as from 7 to 21 Mb on LG VII; Fig 2; Fig 6; Fig S4).

**Table 3.** Genome-wide averages of gene and haplotype diversity in the uplift island populations. Diversity is lower in the potentially younger pond populations on Danger Island and in southeast Montague Island.

| Population | Gene Diversity | Haplotype diversity |
| --- | --- | --- |
| MiFW1 | 0.2771 | 0.395 |
| MiFW2 | 0.3106 | 0.4408 |
| MoSW | 0.3071 | 0.4367 |
| MoSE | 0.2136 | 0.2971 |
| Da04 | 0.2036 | 0.2844 |

**Fig 6.**
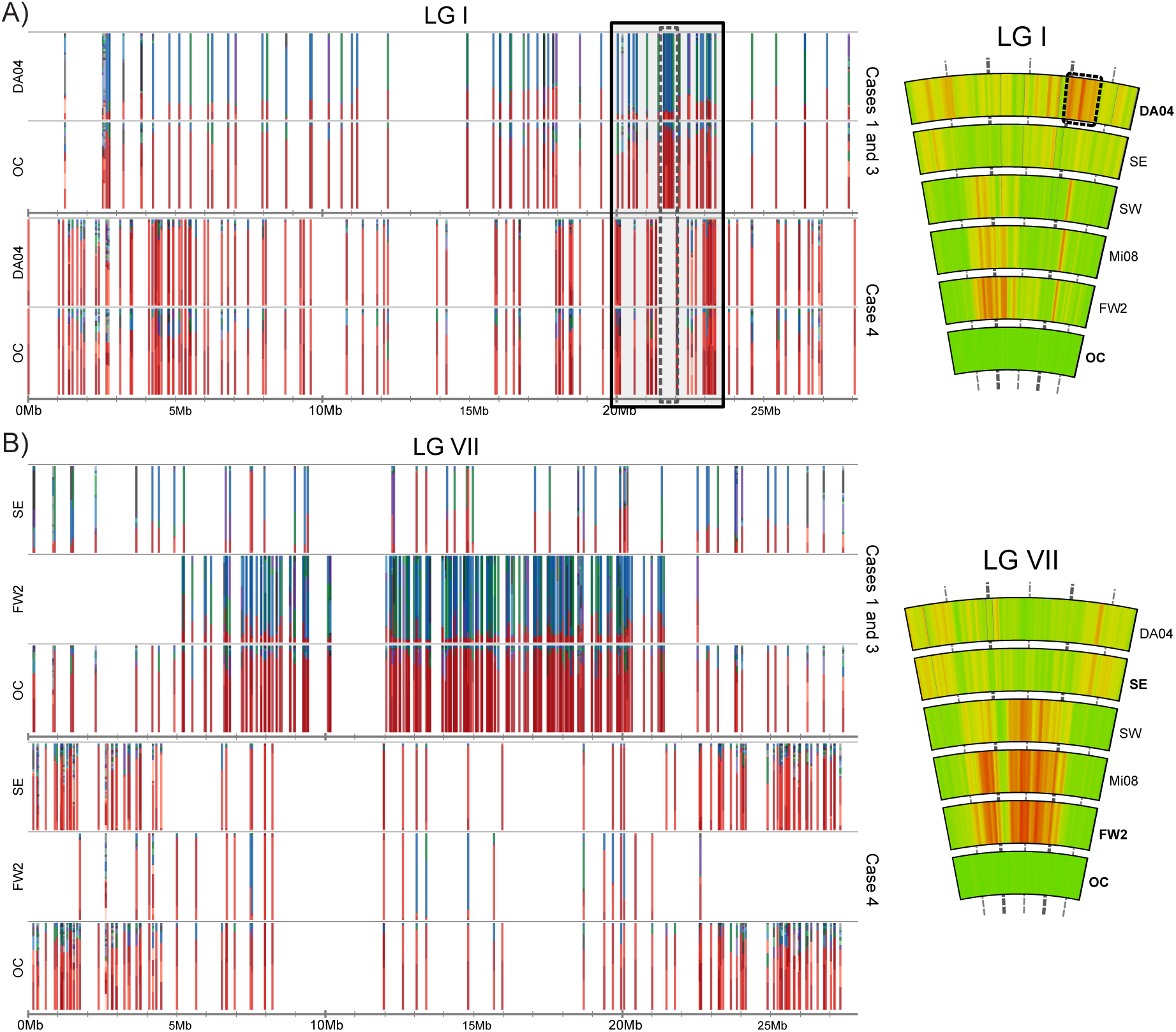
Changes in haplotype frequency reveal ongoing processes in the youngest freshwater populations. The set of haplo types at each locus (where FST´ ≥ 0.2) is represented as a stacked bar chart of haplotype frequencies. Red haplotypes correspond to marine majority haplotypes and blue to fresh water. A) In the Danger Is. FW pond, a LG I region of elevated FST´ (relative to OC) encompassing a known inversion polymorphism shows a strong pattern of FW-OC alternative haplotypes (Case 1 and Case 3 loci) flanked by regions with increased density of loci that are depleted in haplotype diversity relative to OC (i.e., loci classed as Case 4).B) In the southeast Montague ponds, LG VII lacks the high FST´ typically seen across the middle of the chromosome in otherFW-OC comparisons, and instead there is elevated FST´ at the chromosome ends. This inverse pattern is generated by the sparsity of loci with FW-OC alternative haplotypes (Case 1 and Case 3 loci) but an increase in the number loci at which the ocean is more haplotype-rich (Case 4). A clear red-to-blue shift of haplotypes can be seen between marine and FW2 loci on LGVII. For comparison, the FST´ heat-map plots to the right are as in Fig 2.

Danger Island presents another qualitatively distinct scenario from the other freshwater populations in this study; only a fraction of the parallel pattern of freshwater-marine divergence - a single F_ST_´ peak on LG I – is prominent in Da04. This segment encompasses a known inversion whose orientation has been shown to differ consistently between freshwater and marine genomes (62). Relative to OC, Da04 has diffusely elevated F_ST_´ due to genome-wide reduction in haplotype diversity (Fig 2; Fig S3; Table 3), but F_ST_´ crescendos at the LG I inversion. Unlike in the other freshwater groups that share this F_ST_´ peak, however, elevated F_ST_´ relative to OC extends to the inversion’s flanking regions in Da04 (Fig 2; Fig 6). We find that two types of loci primarily occupy the inversion and closely linked regions. In the inversion itself, there is a concentration of loci that have increased in alternative FW haplotypes, and flanking these are loci depleted for common marine haplotypes (Fig 6; Fig S5). Together, these could signal a recent or ongoing selective sweep in Da04.

### Alternative freshwater majority haplotypes are readily detectable at thousands of loci in the marine population

At loci with F_ST_´≥ 0.2, most haplotypes that are in the majority in fresh water can be found in the sea, though they are typically rare (frequency <u>≤</u> 10%) (Table 2; Table S3). For example, 81% of alternative majority haplotypes identified in MiFW1 and 68% of those identified in MiFW2 are found in the marine comparators (131 marine fish from Mi17, Mi23, and Da02) (Table S3). Marine populations that were sampled at great geographic distances from one another - at Middleton Island, at Danger Island (> 100 km mainly across open ocean from Middleton) and even as far as the southern Oregon coast (> 2500 km away, along a coastal route) – carry many of the freshwater majority haplotypes identified on Middleton Is. (Fig 7).

**Fig 7.**
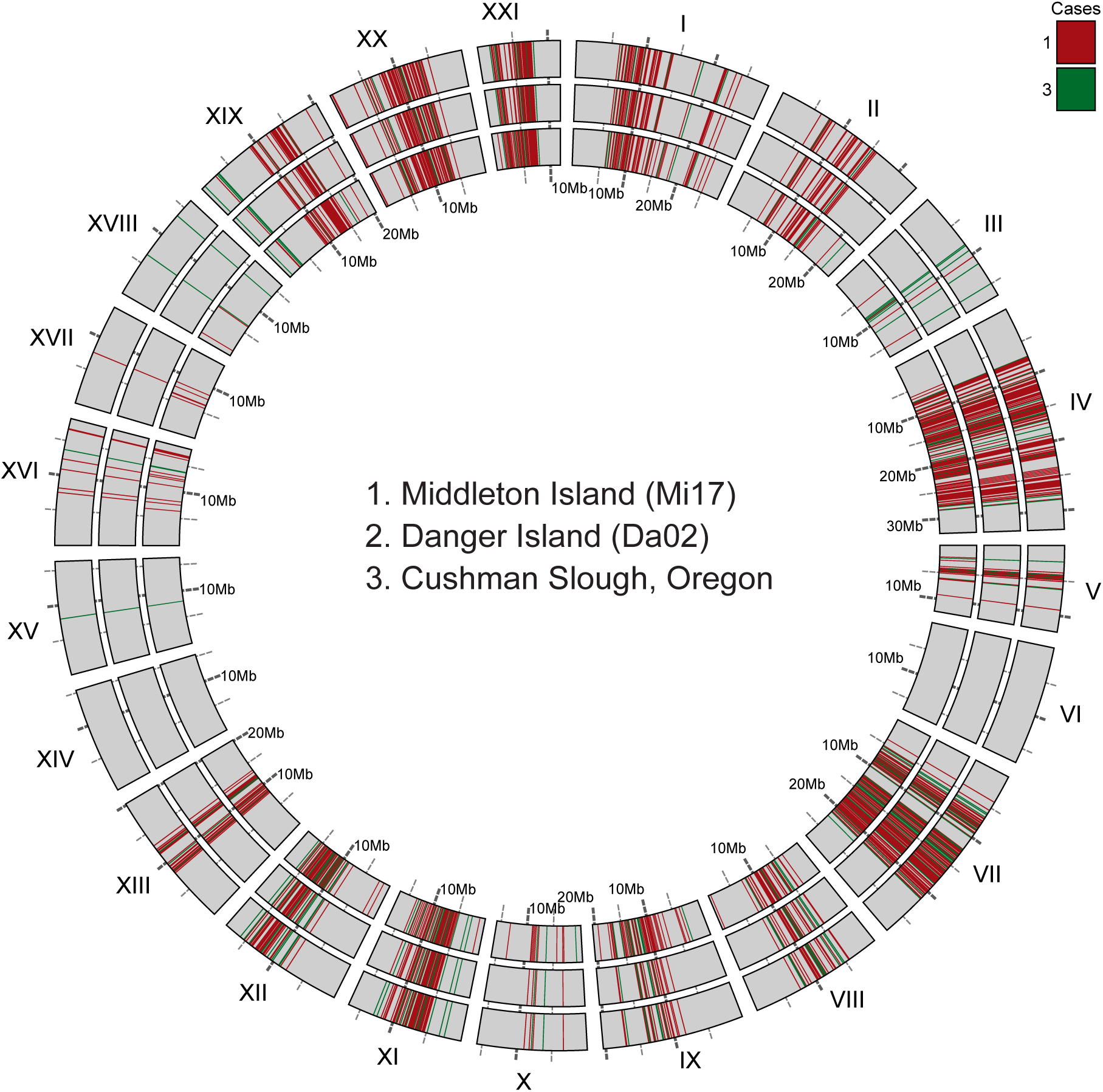
Marine stickleback carry rare haplotypes that are found at high frequency in freshwater fish. Plotted for three geographically distant marine populations are all RAD loci that fit two criteria: the locus falls into Case 1 or Case 3 (as defined in Fig 5) in a comparison between the marine population and a Middleton Island freshwater (Mi11), and at the same locus at least one marine fish carries a FW majority haplotype that is alternative to majority haplotype/s in the sea. Rings are numbered from the center out. Loci are colored by which Case they belong to.

For freshwater haplotypes identified in the island marine populations, the median and mean allele frequency spectra (AFS) at those loci were nearly identical across the three populations (Fig 8; Table S4; Fig S6). In fact, linear models of the pairwise comparison of freshwater allele frequencies across each of the three marine populations resulted in R2 values of 0.69, 0.65 and 0.83 for the Da02:Mi17, Da02:Mi23 and Mi17:Mi23 regressions, respectively. Thus, even between marine populations on two different islands nearly 70% of the variance in allele frequencies of one population are predicted by the other population, and between Mi17 and Mi23 on different coasts of the same island the congruence goes up to nearly 85%. This similarity of freshwater allele frequencies across marine populations holds regardless of whether they were calculated for freshwater majority haplotype sets defined by freshwater populations that are geographically nearby or distant from each marine population (Fig 8; Fig S6; Fig S7). Although the AFS of the entire genome was similar across populations, there was appreciable variation among chromosomes within populations with regards to median and mean freshwater allele frequencies as well as AFS (Table S4; Fig S8), as might be expected given different patterns of divergence across the chromosomes.

**Fig 8.**
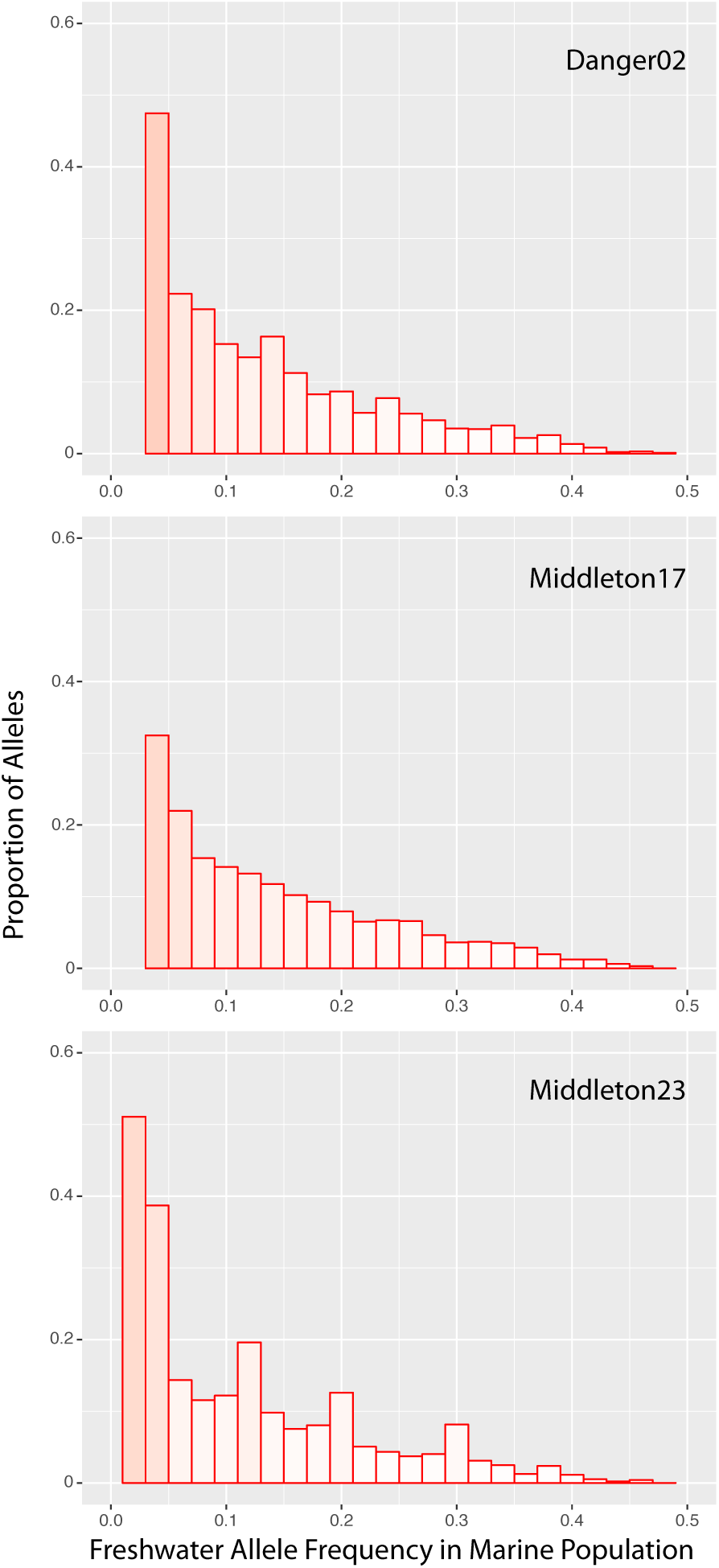
Three marine populations have similar allele frequency spectra of FW majority haplotypes. Distributions of allele frequencies across all Case 1 through 3 loci (as described previously) in comparisons between each marine population and the MiFW1 population. Allele frequency histograms are presented as 2% bins.over 46% of the Case 1-3 freshwater majority haplotypes identified from LG VII, but carries less than 7% of those identified from LG IV, and a Da02 fish harbors over 47% of the LG XII freshwater haplotypes but less than 8% of those from LG VII (Table 4).

The same allele frequencies can be generated if many marine fish each carry a few freshwater haplotypes or if a few marine fish carry a majority or “jackpot” numbers of such haplotypes. To test whether freshwater haplotypes are primarily carried by only a fraction of marine individuals, and to assess the patterns of freshwater haplotypes across the genomes of marine fish, we examined haplotype states at all Case 1-3 loci that carry alternative freshwater majority haplotypes from each marine fish. For example, most of the marine fish in the OC group carried fewer than 10% of these freshwater haplotypes, most often at heterozygous loci, but some fish had much larger numbers of them (Fig S9). Loci bearing freshwater haplotypes are scattered diffusely among linkage groups in most individual marine fish, but in the fish that carry them at the highest proportions, these haplotypes often reside at loci densely concentrated on various subsets of the freshwater-oceanic most divergent linkage groups. For example, one Mi17 individual carries over 46% of the Case 1-3 freshwater majority haplotypes identified from LG VII, but carries less than 7% of those identified from LG IV, and a Da02 fish harbors over 47% of the LG XII freshwater haplotypes but less than 8% of those from LG VII (Table 4).

**Table 4.**
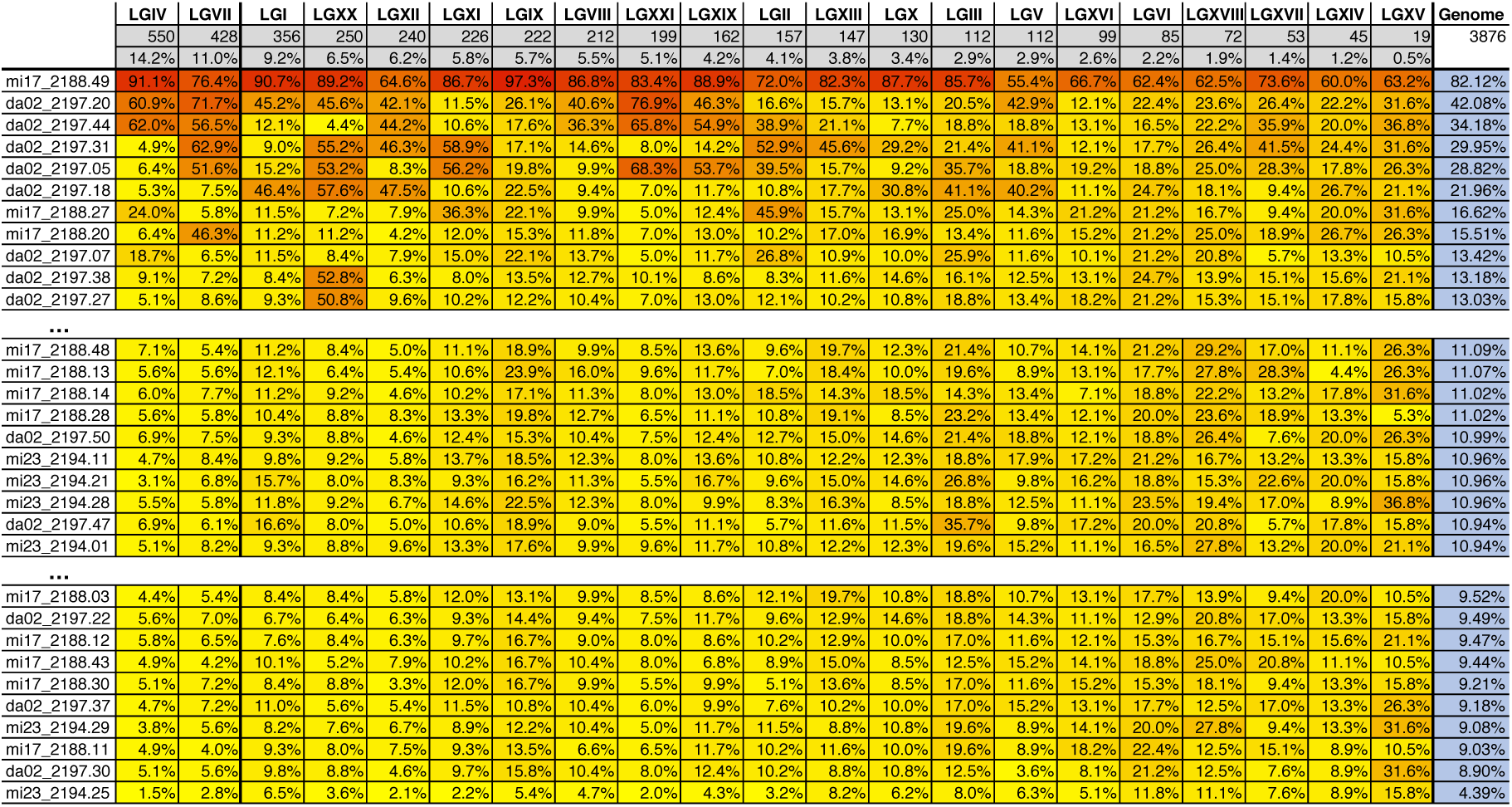
Alternative FW majority haplotypes are carried in a variety of patterns by individual marine fish. Out of the total number of possible FW majority haplotypes identified at Case 1, 2 and 3 loci in the MiFW1 versus OC comparison, the proportion that each marine fish harbors is listed for each linkage group. The total number of possible FW haplotypes is listed first for each chromosome, which are ordered left to right from the highest to lowest number of these loci. Some chromosomes, such as LG XV, have relatively few loci with high FW-OC divergence, so there are few possible FW haplotypes on them. Others, such as LG IV and VII have very large numbers of these loci. Cells are heat map colored to highlight where individuals harbor a higher (red) or lower (yellow) proportion of the possible FW haplotypes on a given chromosome. Individuals are ranked from those that carry the most to the fewest FW haplotypes, and of the 131 marine fish sampled, representative blocks of fish at the top, middle and bottom of this range are shown.

## Discussion

### Heterogeneous genomic divergence occurs in just decades despite gene flow

The heterogeneous divergence across the stickleback genome that we documented in these very young populations is unlikely to have occurred simply by neutral demographic processes or by the action of background selection in genomic regions of reduced recombination (63-66). The stickleback genome is nearly dichotomous in these newly formed populations, with clustered regions of highly significant genetic differentiation on a background of no divergence. Furthermore, this genomic mosaic is replicated across independently derived populations, which is not expected under neutral processes or background selection alone. The most plausible explanation is that strong, alternative selection between the marine and freshwater habitats has rapidly changed allele frequencies in only a subset of genomic regions.

These so-called “genomic islands” (67-69) may be created by the action of local selection that sorts ancestral variation either in the absence of contemporary gene flow,or with continuing gene flow via a process termed ‘divergence hitchhiking’ (9, 70, 71). Because of the very recent origin of the freshwater populations on the uplift islands, discriminating these two processes is difficult. However, some characteristics of these populations argue for the opportunity for gene flow during divergence since the 1964 earthquake. Marked reduction in body size and bony lateral armor plates is typical of freshwater stickleback (38), and we documented previously that many uplift ponds on Middleton house only small fish with minimal armor (35). Several low-lying ponds still allow influx of marine stickleback, however, and harbor both low-armored and well-armored fish as well as some phenotypic and genetic intermediates (35). Even here, despite cohabitation and evidence of hybridization, the same divergent genome segments seen in older, more isolated freshwater populations (46, 72) manifest when sympatric marine and freshwater morphological ecotypes are compared against each other (seen in F_ST_´ in Fig 2). In addition, these regions of divergence align with defined blocks of linkage disequilibrium (LD) that stretch over megabases and mirror the LD pattern of an *in silico* “mock sympatric population” in which data from allopatric marine and freshwater samples are artificially combined (Fig S10). The LD pattern implies that such ponds are zones of contact between alternatively adapted fish.

### Strong alternative and divergent selection in both marine and freshwater habitats

Historically, a central (but not exclusive; see (73, 74)) focus of stickleback phenotypic and population genomic studies has been on the action of natural selection in freshwater habitats as marine fish colonize them. However, the present study underscores the action of similarly strong divergent selection in marine habitats. The reuse of ancient haplotypes can be seen in comparisons among the independently colonized islands. In fact, we observed as much as 99% of freshwater majority haplotypes shared between freshwater populations are at RAD loci where marine and freshwater populations have strongly alternative haplotypes. Although it has been predicted that a pool of freshwater-adapted haplotypes must be rare but present in and dispersed by the marine population (e.g., (26, 75)), such haplotypes had so far not been quantified with a genome-wide approach. In a sample of only 131 phenotypically marine stickleback collected along the shores of the uplifted islands, we found a large proportion of freshwater majority haplotypes that had been identified at divergent loci in island pond populations (Fig 7). As would be predicted if many freshwater-adapted haplotypes are selected against in the sea, we found them to be rare and most often carried heterozygously in marine stickleback (Fig S9). Nonetheless, the allele frequency spectrum of freshwater majority haplotypes borne by the marine populations (Fig 8) argues that continuous and substantial gene flow from myriad freshwater habitats maintains appreciable levels of these genotypes despite selection against them in the sea.

Together, our data and other recent studies in stickleback argue for a view of the stickleback species as a metapopulation, with strong divergent selection occurring in both marine and freshwater habitats despite bi-directional gene flow between them. This view contrasts with a more traditional, ‘raceme’ or ‘bottle-brush’ concept of stickleback, with a core of marine populations that harbor the majority of genetic diversity in this species, while freshwater populations are the bottle-brush ‘whiskers’, ephemeral in nature (so-called “evolutionary dead-ends“). In this asymmetric model, the freshwater populations draw on the genetic diversity stored in the large marine population, but themselves wink out of existence having had little influence on the genetic composition of the species as a whole. In contrast, our data argue strongly that both population types serve as reservoirs for maintaining genetic diversity through sustained bi-directional gene flow that is sorted by divergent selection in both habitats, in other words, through a long-term dynamic of migration-selection balance. Although the relative impact of any particular freshwater population may be slight, the aggregated millions of freshwater populations could have a significant impact on the pool of standing genetic variation in marine populations. This is particularly likely when, as has been documented, haplotypes that are identical by decent are selected repeatedly in independently derived freshwater populations.

### A large proportion of the stickleback genome is affected by selection during divergence

By comparing patterns of freshwater-marine divergence in the young island populations against post-glacial mainland populations, we find that as much as 15% of the genome in common is affected by alternative adaptation to ponds and lakes versus the sea (Table 1; Table S2; Fig S2). By comparison, Jones *et al.* (62) sequenced 20 genomes in a geographically broad survey of marine and freshwater stickleback, and found that less than 0.5% of the genome fell in a phylogenetic dichotomy of freshwater and marine alleles. The findings from these two studies are of course strikingly divergent, and several factors may explain this 30-fold difference between them in the inferred proportion of the genome involved in alternative adaptation. Importantly, differences in the methodologies employed between the studies in terms of geographic breadth versus population genomic depth of coverage suggest that the findings are likely compatible and provide different and complementary information on divergence of stickleback between marine and freshwater habitats. A specific subset of selected loci – those that are both presumably ancient and house strictly alternative haplotypes - can be revealed in a broad geographic survey of few individuals, whereas polygenic selection may involve more subtle changes in allele frequencies at multiple loci (76, 77) that can only be detected with deeper population genomic sampling. Similarly, the immediate effects of linked selection can be revealed by this kind of sampling, particularly in very young populations such as on these uplifted islands.

While we might expect to see broader regions of divergence in these young freshwater ponds because of the effects of linkage (Table 1), others investigating stickleback populations that could be of similar age to those we studied here reported much less total divergence across the genome. Terekhanova *et al.* (42) studied marine-freshwater genomic divergence along the shores of the White Sea (Russia) in populations that were of known or inferred age ranging from several decades to several centuries. Those of known age — in two flooded rock quarries — were artificially seeded with only two or forty individual stickleback from both marine and freshwater habitats, making it difficult to assess the relevance of the findings to the dynamics of natural populations. They also focused on four freshwater populations that were naturally formed but of less certain independence and age, inferred from projections of sea level rate of change. The proportion of the genome estimated to be affected by marine-freshwater divergence in these Russian populations is far less than what we report for young populations on the uplift islands or even for much older Alaskan populations (Fig S11; Table 1; Table S2). Regional differences between the White Sea and Alaskan stickleback could account for these broad differences. Alternatively, the small biological sample sizes from the White Sea populations, and subsequent analytical approaches that this under-sampling necessitated (42), could have led to a significant underestimate of the extent of linked selection there.

In contrast, because of the high level of biological sampling at the level of individuals and populations, we provide clear evidence that at least a quarter of the stickleback genome is initially affected by the action of strong divergent natural selection, and that this effect is replicated independently across populations within and among islands. A key, novel finding that emerges from such a sufficiently powered population genomic study is an unexpected change in haplotype diversities in each habitat. Regions of reduced haplotype diversity in the marine populations correspond to most of the genomic segments elevated in F_ST_´ (Fig 4), a pattern that may have gone unrecognized in previous oceanic-freshwater comparisons because of the primary use of SNPs (43 72) or sampling that does not allow accurate diversity estimation within populations (42, 48). For example, Terekhanova *et al.* (42) report that nucleotide diversity in the ocean is higher than in fresh water within regions of divergence. This important difference may be due to sampling (Fig S11) or to uncertainties in using nucleotide diversity as a proxy for haplotype diversity. Regions of reduced absolute and relative haplotype diversity in the sea highlights an under-appreciated likelihood of strong directional selection in both habitats, not just in fresh water.

That haplotype diversity is relatively higher in freshwater fish across the regions of divergence perhaps runs counter to an intuition that hard selective sweeps could reduce haplotype diversity in nascent freshwater populations relative to genetically diverse holarctic marine populations. One potential explanation for elevated diversity in these young populations is that freshwater-adapted haplotypes have not completely displaced marine haplotypes, as they are in a transitory state due to limited time since founding or they are in a stable equilibrium due to lack of isolation from the sea. A high magnitude of F_ST_´ divergence, however, can also manifest through a relative increase in freshwater haplotype diversity *via* selection of an ancient allele embedded in a small number of syntenic genotypes, a process that would transiently increase the relative haplotype diversity in neighboring genomic regions in freshwater populations (78). The majority of selected genomic regions may comprise such old alleles - for example a region highlighted in the recurrent evolution of armor loss (55, 75, 79) *-* which drive the broadly parallel trait changes in freshwater fish, but these alleles exist on diverse genomic backgrounds.

An important extension of our findings is that linked selection across these large regions of divergence may drive significant correlated phenotypic responses in both maintenance of marine-adapted traits in the sea and alteration of traits following freshwater colonization, even if the overall proportion of genetic variants that are subject to direct selection in common across populations is relatively small. Our observation supports the role of extended linked selection affecting a large proportion of the stickleback genome, and accords with a documented pattern of greater phenotypic variation among geographically localized freshwater populations relative to the globally distributed marine population (36). This broader phenotypic range has been attributed to the diversity of limnological habitats (36). Findings from these Alaskan island populations suggest an additional hypothesis: that divergent phenotypic responses could result from direct selection on a subset of alleles and indirect selection on a diverse array of linked variants, leading to a correlated response in both mean and variance of traits. We studied only a dozen populations on three islands, but the geographic region uplifted in 1964 likely led to many additional freshwater stickleback populations that could provide a key resource for future studies.

### The transporter hypothesis and the size and ordering of genomic changes in the first few generations after colonization

Migration-selection balance and the reuse of ancient haplotypes underlie the popularly cited “Transporter Hypothesis” to interpret parallel evolution in threespine stickleback (26). These mechanisms predict that the freshwater adapted genome should enter new habitats piecemeal from marine colonizers over a period of time and that populations in the earliest stages of colonization might therefore initially have captured only a subset of these ancient haplotypes. Though capped at a maximum age by the 1964 earthquake, the island populations studied here probably span a range of younger ages. Montague’s SE ponds that are perched on a narrow peninsula less than two meters above sea level (Fig S12), and similarly, the only freshwater pond (Da04) on Danger Island sits only one meter above sea level. These ponds might house the youngest freshwater populations in this analysis, as they are vulnerable to periodic inundation from the sea, which could cause osmotic stress, local extinction of adapting freshwater stickleback, or an influx of marine stickleback leading to recolonization or genetic introgression. These populations show only a subset of the pattern of divergence seen in the other freshwater groups(e.g., (51) and this study). For instance, an inversion-spanning region on LG I (62) alone stands out in Da04 (Fig 2; Fig 6). While Middleton populations and SW Montague have a sharp peak of F_ST_ coinciding with this small inversion, in Da04 the inversion sits in a broader window of divergence that includes reduced haplotype diversity, the clear signature of an ongoing selective sweep. The sparser pattern of genomic divergence in these precarious habitats may reflect initial random sampling of freshwater-adapted genotypes from the sea, and could also imply that these subsets of loci are sufficient for initial persistence of freshwater pioneers. Along those lines, others have remarked on the location at the LG I inversion of *atp1a1*, a gene encoding a sodium-potassium ATPase important for osmoregulation in fishes (80-82).

Though fewer chromosomes show divergence from marine fish in MoSE, it nonetheless has elevated F_ST_´ across the full length of the massive region of parallel divergence on LG IV (Fig 2). This is perhaps a testament to a low recombination rate across this part of the chromosome (65), making it more likely for LG IV haplotypes alternative to majority marine haplotypes to enter new ponds as a unit. If true, this genomic architecture should speed the time that the parallel pattern of a diverged freshwater-adapted genome can be reassembled, from what might be a relatively small number of large genomic segments.

Most of the alternative freshwater majority haplotypes we identified from MiFW1, MiFW2 and MoSW were also present in marine stickleback, where in most individuals they occur in low numbers across the genome. However, a subset of the marine fish in our sample carried freshwater haplotypes densely on some linkage groups (Table 4). These individuals could be early generation hybrids between freshwater and oceanic stickleback. Some of the freshwater haplotypes may also be in regions that resist recombination, such as those that lie in inversions or across large parts of LG IV and LG VII (65). Discovery of these patterns of freshwater majority haplotypes in a relatively small sample of fish from the sea suggests that the rapid time of adaptation to new freshwater habitats could hinge on “jackpot” carriers, and that a freshwater adapted genome might effectively be reassembled by migrants in new habitats one or more chromosomes at a time. Together, our findings provide a potential answer to the question of how enough low frequency standing genetic variation can be reconstituted in small freshwater ponds even with few initial stickleback colonists: luck of the draw. A small number of migrants may in total carry enough chromosome-scale freshwater haplotypes to colonize newly opened habitats. Even if the initial colonists do not have an entire set of variants, subsequent migrants may quickly complete the winning hand.

### General lessons about migration-selection balance and the genomic architecture of rapid adaptation

Striking genomic patterns of strong differentiation on a background of virtually no divergence argue that changes during adaptation to fresh water are established promptly, and that mainland populations in much older lakes could have experienced most of their present-day genomic shifts, and associated phenotypic evolution, in just their first decades after colonization. The speed with which freshwater and marine populations diverge across a large number of genomic spans - on the order of decades - could be an outcome of the metapopulation structure particular to stickleback. On the other hand, the stickleback radiation may provide more general evolutionary genomic lessons because adaptation from standing genetic variation on such a short time frame may be common in the wild. This could be particularly true in spatially structured species that experience variable selection and gene flow that maintains ancient diversity, a scenario we argue is more likely to be the rule than the exception in natural populations of most species, as has been documented, for example, in the study of Darwin’s finches (83). If common, then for many species standing genetic variation maintained and shaped by molecular evolutionary processes operating over thousands or millions of generations may permit population genomic changes and organismal evolution to new environments over just decades. Fully understanding rapid adaptive dynamics may often require a deep molecular evolutionary perspective of the haplotypes’ changing frequencies.

The long-term association of the stickleback species with both oceanic and freshwater habitats leads to an intriguing hypothesis. Although each bout of adaptation may only require decades, the genomic regions underlying the change are a much older balanced polymorphism. If so, the molecular evolutionary history of locally adapted genomic regions across the metapopulation might actually be quite deep. Alternatively-adapted freshwater and marine haplotypes in genomic regions with lower recombination rates (65) may be thousands or millions of years old (75) and lead to increased absolute divergence even in the absence of contemporary gene flow. In fact, longer read RAD-seq data from our laboratory on many fewer individuals align with this hypothesis. Coalescent analysis of these Sanger length sequences sampled across the genome demonstrates that in many of the genomic regions with the most significant relative divergence of allele frequencies, the haplotypes also show more ancient evolutionary divergence that is millions of years old (84). Therefore, a deep evolutionary history underlies the ability of stickleback to adapt so rapidly. Similar analyses of other species with populations experiencing divergent selection and gene flow could show whether this phenomenon is unique to stickleback, or a common theme in nature.

## Materials and methods

### Biological collections

Freshwater ponds and marine sites for stickleback collection (Fig S1) were chosen on the basis of maps and aerial imagery created before and after 1964. Fish were trapped during the summers of 2005 (Montague and Middleton Islands), 2010 (Danger and Middleton Islands) and 2011 (Montague and Middleton Islands), and processed as described in Lescak *et al.* (35). All research was approved by the IACUCs of the University of Alaska and the University of Oregon. Fish were collected under Alaska Department of Fish and Game permits SF-2005-020, SF-2010-029, and SF-2011-153, as well as Oregon Department of Fish and Wildlife scientific taking permits OR2007_3495 and 13920.

### Restriction site Associated DNA sequencing library preparation and sequence analysis

RAD-seq libraries for 1057 fish were created as described in Lescak *et al.* (35) using endonuclease *SbfI*, and sequenced to 101 nucleotides (including a 6 nt inline barcode) on an Illumina HiSeq 2000 platform (Table S5). In addition to these data, RAD sequences from an additional 98 fish from Cushman Slough, Oregon (USA), which were first published in Catchen *et al.* (85), were included in analyses for a total of 1155 fish. Individual fish were represented by an average of 1,265,744 ±13,864 SE sequences each, of which 1,225,729 ±13,744 SE sequences (97%) were aligned to the reference genome, and these produced an average of 43,174 ±165.7 SE loci with an average depth of coverage of 27x using the software *Stacks* (86-88), version 1.46 (Table S6). Raw sequence data were demultiplexed according to barcode and filtered for quality using the process_radtags module (-`c`, -`q`, and -`r` parameters) in *Stacks*. Cleaned reads were aligned to the stickleback reference genome (version BROADS1, Ensembl release 86) using the mem module from the BWA aligner (89) with default settings. *Stacks* were assembled, SNPs were called and a population-level catalog was constructed by executing the pstacks (-`m` 3), cstacks (–`aligned`), and sstacks (–`aligned`) modules. Population-level corrections were made to the data by running rxstacks (–`model_type bounded`, –`bound_high` 0.1, –`prune_haplo`, and –`conf_lim` 0.25 parameters), and then rebuilding and matching to the catalog with cstacks (–`aligned`) and sstacks (–`aligned`). Final filtering was done with the populations module, as well as the calculation of population genetic statistics and associated kernel smoothing.

Our full data set comprises 48,307 RAD markers encompassing a total of 266,902 SNPs. We applied two general filtering steps for most analyses, one when considering populations individually, one when considering Structure-defined population clusters. In the former, for a locus to be included in downstream analyses we required it to be present in at least 20 populations (-`p` 20 parameter) and in 75% of individuals of each population (-`r` 0.75), and we applied a minor allele frequency (MAF) filter of 1% (–`maf` 0.01). This resulted in 29,911 RAD markers and 74,556 SNPs. In the latter analysis, a locus had to be present in 9 populations and in 75% of individuals, with a MAF of 1%. This resulted in 30,285 RAD markers and 75,476 SNPs. Sequence data are available in the NCBI Short Read Archive (http://www.ncbi.nlm.nih.gov/sra) under accession XXXXX.

To compare the Middleton Island samples against those found in Cook Inlet previously, we downloaded sequences used in Hohenlohe *et al.* (51) from the NCBI Sequence Read Archive using accession SRA010788.9. In addition, we added 54 more recently sequenced individuals from Rabbit Slough and 24 from High Ridge Lake. As these data stem from sequencing runs using different versions of Illumina technology, we retained 94 individuals that had read lengths of at least 49bp and an average depth of coverage of 10x (Table S7). These data were aligned to the stickleback reference genome and processed with *Stacks*, in the same manner as the Middleton Island data. We grouped the individuals into freshwater and marine clusters, required loci to be present in both clusters and at a frequency of 75% among individuals in either separate cluster, and applied a MAF of 6%.

### Statistical approach

The populations that were shown to be united into genetic groupings by Structure analysis in Lescak *et al.* (35) were treated together here as groups rather than as single populations. Also, the three geographically proximate Cook Inlet freshwater populations sampled in Hohenlohe *et al.* (51) are here treated as a group to increase sampling robustness for a regional comparison. Latitude and longitude, respectively, for these three populations are 61.6 and -149.75 (Bear Paw Lake), 61.72 and -150.12 (Boot Lake), and 61.93 and -150.97 (Mud Lake). See Fig S1 for information on island sampling locations and population groupings. Population genetic statistics were calculated using the populations program in *Stacks* (86-88). Calculations of haplotype diversity, ϕST and F_ST_´ are based on the state of each ∼100bp RAD locus instead of individual SNPs. The combination of fixed and variable nucleotides that make up each locus are considered a set of haplotypes at the locus. While SNP-based statistics require bi-allelic nucleotide positions, the haplotype-based statistics can have multiple haplotypes per locus, with a distance measured between haplotypes, providing a richer set of states. Genome-wide circle plots (*aka*, Large Hadron Collider Plots) were generated by a Python program, lhc_plot.py, which is distributed with the *Stacks* package. Population genetic statistics were kernel smoothed along the genome using default smoothing parameters in *Stacks* software.

### Calculating linkage disequilibrium (LD)

All SNPs output from *Stacks* were imported into Beagle (version 3.3.2) to phase SNPs across each chromosome of each population. We then transferred the resulting Beagle output into the phasedstacks program of *Stacks* to calculate D´, the measure of LD (90, 91), and values of D´ were plotted as heat maps in Gnuplot 4.6 patchlevel 4 (http://gnuplot.info).

### Calculating Gene and Haplotype Diversity

We calculated two separate diversity measures for each RAD locus in each population. A RAD locus contains one or more haplotypes, which are the physically phased combination of SNPs present at that locus across the population. The first measure, Gene Diversity (90, 91), can be thought of as the probability that two randomly chosen haplotypes at that locus are different. Given the number of samples *n*, and the *ith* haplotype *p*, it is calculated as:

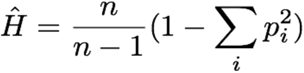

The second measure, Haplotype Diversity (90, 91), is similar to Gene Diversity, but it is scaled by the substitution distance, *d*, between the two randomly chosen haplotypes:

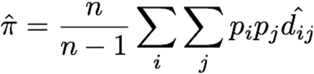

Haplotype diversity measures were kernel smoothed along the genome using default smoothing parameters in *Stacks* software. For haplotype diversity difference, the continuous smoothed values for the marine group were subtracted from those for the freshwater group, and these raw values were plotted on a nonlinear color scale (e.g., as in Fig 3E).

### Segmenting the genome via Hidden Markov Model

We used the General Hidden Markov Model Library (GHMM (92), version 0.9-rc3) to segment the genome using haplotype-based F_ST_´. In order to make the F_ST_´ measures discrete, we binned them into four categories; we took the F_ST_´ measures from LG IV of the FW2 versus marine data set and hierarchically clustered them in R (R Core Team, 2013). Using the Python bindings for GHMM, we designed a two-state HMM representing *diverged* and *not-diverged.* Initial emission and transition parameters were defined using F_ST_´ values in regions of marked divergence between freshwater and oceanic populations on LG IV and in regions of nearly zero divergence on LG XV, respectively. For each pairwise comparison (FW2 vs. marine, FW1 vs. marine, SW vs. marine, and Cook Inlet vs. marine) the HMM was trained on LG IV (diverged) and LG XV (not diverged) by executing the Baum-Welch algorithm of the GHMM library on the FW2 vs. marine F_ST_´ values. After training, each linkage group was segmented into the *diverged* and *not-diverged* states using a custom program segment.py. Finally, we compared the set of states for each of the four data sets (FW2, FW1, SW, CI vs. marine) to find the diverged regions in common using our shared_segments.py program. Plots of the HMM data were generated using Gnuplot 4.6 patchlevel 4 (Fig 3; Fig S2).

### Defining cases from haplotype data

Classification of cases was done using custom Python scripts. For each locus in an analysis, we summed the haplotypes in order from the haplotype with the largest frequency in a population to the smallest. The set of ordered haplotypes that occupied at least 60% of the haplotypes at that locus in that population were considered *majority haplotypes*. Loci were classified as *Case 1* when one or more majority haplotype/s occurred in a marine population, but the corresponding freshwater population had a single, alternative majority haplotype (i.e., one that is *not* a majority haplotype in the marine population). Loci were classified as *Case 2* when the marine population had one or more majority haplotype/s at a locus and the freshwater population had one alternative majority haplotype, and one or more haplotype/s also in the majority in the marine population. Loci were classified as *Case 3* when the marine population had one or more majority haplotype/s at a locus, but the corresponding freshwater population had two or more alternative majority haplotypes and no majority haplotypes in common with the marine population. Loci were classified as *Case 4* when the freshwater population had only a subset of majority haplotypes in the marine population. Circular genomic plots of cases were created with the lhcp_plot.py script while the linear plots of cases (e.g., Fig 6) were created with a custom Python script.

## Acknowledgements

We thank M. S. Christy, S. A. Hatch, B. Lohman, V. M. Padula, L. Smayda, and K. Walton for assistance with fieldwork logistics and fish collection, as well as T. Wilson and M. Currey for technical help. We also thank P. Hohenlohe for discussions during early stages of the development of the project. We thank members of the W.A.C., P. C.Phillips, and M. A. Streisfeld Laboratories at the University of Oregon, for discussions and critical comments.

